# The unique face of anxious depression: Increased sustained threat circuitry response during fear acquisition

**DOI:** 10.1101/2023.10.17.562565

**Authors:** Tate Poplin, Maria Ironside, Rayus Kuplicki, Robin L. Aupperle, Salvador M. Guinjoan, Sahib S. Khalsa, Jennifer L. Stewart, Teresa A. Victor, Martin P. Paulus, Namik Kirlic

## Abstract

**Background:** Sensitivity to threat with dysregulation of fear learning is thought to contribute to the development of psychiatric disorders, including anxiety disorders (AD) and major depressive disorder (MDD). However, fewer studies have examined fear learning in MDD than in AD. Nearly half of individuals with MDD have an AD and the comorbid diagnosis has worse outcomes. The current study used propensity matching to examine the hypothesis that AD+MDD shows greater neural correlates of fear learning than MDD, suggesting that the co-occurrence of AD+MDD is exemplified by exaggerated defense related processes.

**Methods:** 195 individuals with MDD (N = 65) or AD+MDD (N=130) were recruited from the community and completed multi-level assessments, including a Pavlovian fear learning task during functional imaging.

**Results:** MDD and AD+MDD showed significantly different patterns of activation for [CSplus– CSminus] in the medial amygdala (ηp^2^=0.009), anterior insula (ηp^2^=0.01), dorsolateral prefrontal cortex (ηp^2^=0.002), dorsal anterior cingulate cortex (ηp^2^=0.01), mid-cingulate cortex (ηp^2^=0.01) and posterior cingulate cortex (ηp^2^=0.02). These differences were driven by greater activation to the CS+ in late conditioning phases in ADD+MDD relative to MDD.

**Conclusions:** AD+MDD showed a pattern of increased sustained activation in regions identified with fear learning. Effects were consistently driven by the threat condition, further suggesting fear signaling as the emergent target process. Differences emerged in regions associated with salience processing, attentional orienting/conflict, and self-relevant processing. These findings help to elucidate the fear signaling mechanisms involved in the pathophysiology of comorbid anxiety and depression, thereby highlighting promising treatment targets for this prevalent treatment group.

## Introduction

Major depressive disorder (MDD) co-occurring with an anxiety disorder (AD), such as generalized anxiety disorder (GAD), social phobia, panic disorder, or simple phobia (collectively referred to as AD+MDD), represents the most prevalent form of presentation in mental health care, accounting for nearly 50% of MDD cases (1). Yet, specific studies of this comorbidity are rare. Gaining insight into the fundamental process dysfunctions is paramount for identification of distinct subtypes of MDD with underlying mechanisms that could be more receptive to targeted interventions, such as neuromodulation of circuits associated with a specific process dysfunction. By categorizing MDD into subtypes based on anxiety comorbidity, the heterogeneity among MDD patients in clinical trials could potentially be reduced, in turn potentially mitigating mixed findings (2).

AD+MDD is linked to increased treatment resistance (3), less favorable treatment results (4), more rapid symptom recurrence (5), and elevated suicidal thoughts (1) compared to MDD without an AD. Despite this more challenging prognosis, engagement with treatment tends to be higher in cases of AD+MDD (1). Standard first line treatments include psychotherapy (e.g., cognitive behavioral therapy) and second-generation selective antidepressants (e.g., selective serotonin reuptake inhibitors). Second line treatments include first-generation mixed target antidepressants (e.g., tricyclics) potentially augmented with atypical antipsychotics and benzodiazepines for AD (6, 7). There has been a move away from the use of benzodiazepines because of the risk for use disorder and unfavorable side effects profiles, which can be increased in AD+MDD (8). Therefore, due to the high incidence rates and unique complexities of AD+MDD such as increased resistance to first-line pharmacological treatments and a heightened likelihood of experiencing adverse drug reactions it is important to conduct in-depth research that aims to identify distinct neurobiological subtypes and elucidate the underlying neural processes. Such research will facilitate the development of personalized intervention strategies, thereby improving the likelihood of achieving favorable treatment outcomes in this particularly challenging patient population.

MDD and AD have broad overlapping neurocognitive process dysfunctions, primarily related to negative affect or general distress (9, 10). However, studies examining potential neurocognitive differences between MDD and AD+MDD are rare, which may be impeding treatment selection and development. One complicating factor is the categorical nature of diagnosis, which has been tackled with approaches that seek to use specificity of process dysfunction to organize and describe phenotypes relating to psychopathology (10, 11). This separates ADs into “fear” and “distress”, disorders, with phobias and panic disorder being associated with “fear” and GAD being associated with “distress”, the latter having more overlap with MDD. However, common across ADs is increased threat related attentional bias or threat sensitivity (12), a maladaptive process associated with heightened awareness and reactivity to threats in one’s environment.

Threat sensitivity, a multidimensional construct encompassing physiological, cognitive, affective, and behavioral responses to aversive stimuli, is a salient feature in both AD and comorbid AD+MDD (13). Threat sensitivity in AD+MDD is associated with dysregulation of stress circuitry (14). Threat responses associated with AD have been shown to engage a corticolimbic circuit including ventromedial prefrontal cortex (vmPFC), insula and amygdala (15, 16). This construct manifests differently through its transdiagnostic facets, namely anxious apprehension, characterized by repetitive negative thinking, and anxious arousal, marked by hypervigilance and hyperarousal of the sympathetic nervous system. Although the neural correlates of these facets are still being mapped, there is emerging evidence for distinct circuitry. Anxious apprehension, associated with increased activity in the dorsolateral prefrontal cortex (DLPFC), vMPFC, hippocampus and left anterior insula (17, 18), may contribute to the cognitive aspects of threat sensitivity in AD+MDD. On the other hand, anxious arousal, linked to increased activity in the amygdala, periaqueductal grey area and right anterior insula (19–21), may be particularly relevant for understanding the physiological dimensions of threat sensitivity in this population.

In comparison, non-anxious MDD often displays a blunting or desensitization of anxious arousal (22, 23), indicating a different position on the threat sensitivity spectrum. This suggests that a nuanced, adaptive level of threat sensitivity likely exists between the blunted responses observed in non-anxious MDD and the heightened sensitivity seen in maladaptive anxiety or AD+MDD. Therefore, lumping AD+MDD and MDD into a single clinical sample for comparative studies with healthy comparisons can be deceiving, as these clinical groups represent opposite ends of the threat sensitivity spectrum. The intricate interplay between MDD and AD, including their diverse neurocognitive processes and a spectrum of threat sensitivity, underscores the need for neuroscience-informed diagnostic approaches and targeted research. Elucidating these relationships using multi-level data (symptoms, physiology, behavior, and neural circuit measures) will facilitate more accurate differentiation and inform treatment selection, thereby mitigating the risk of obscuring essential variations that could significantly impact therapeutic strategies.

Pavlovian fear conditioning is a well-studied phenomenon of associative fear learning, with foundations in preclinical literature (24). Briefly, animals learn an association between a conditioned stimulus (CS, a set of visual, auditory, or olfactory cues) being paired (CS+) or unpaired (CS-) with an unconditioned stimulus (US, an aversive outcome such as electric shock or aversive noise), with CS+ enabling fear learning and CS-enabling safety learning. An early role for the amygdala was established in preclinical models (24) and in the last 30 years this has been explored in humans using neuroimaging. Human findings are mixed, particularly for the amygdala (25), likely due to historically small sample sizes (*n* = 20-50) for neuroimaging, whilst lesion studies were more consistent (26). A recent review (27) proposes a role for the amygdala, anterior insula and cingulate cortex in fear learning and the vmPFC and DLPFC in safety learning. Findings in psychopathology are also mixed, with AD patients showing increased ACC, anterior insula and (less consistently) amygdala (28) activation during fear acquisition, effects not seen in MDD (29). Heightened skin conductance responses during fear acquisition in AD versus blunted responses in MDD were also reported (30). The potentially opposing forces of depression and anxiety on fear acquisition makes the study of comorbid AD+MDD of particular interest.

The current investigation aimed to determine whether AD+MDD shows a unique profile of exaggerated defense related processes (i.e. threat sensitivity) compared to those with MDD alone, measured by threat circuitry (amygdala, anterior insula and cingulate cortex) activation during fear conditioning, using neuroimaging data from a large transdiagnostic sample collected as part of the Tulsa 1000 study (31). The basic approach was to compare a propensity-matched sample of MDD and AD+MDD individuals on neural activation during fear conditioning. We previously showed that those with AD+MDD showed exaggerated threat sensitivity using startle electromyography (32), and interoceptive/nociceptive sensitivity via a breath hold task and a cold pressor task (33) compared to those with non-anxious MDD who showed valence-independent blunting of emotional responses. Here, we hypothesized that those with AD+MDD would show increased neural responses in threat circuitry (amygdala, anterior insula, and cingulate cortex) to fear acquisition (driven by CS+) compared to those with non-anxious MDD, driven by increased threat sensitivity. We will also explore group differences in the activation of regulatory circuitry (DLPFC, vmPFC) during fear acquisition, with the aim to give further insight into the top-down/bottom-up dysfunction underlying threat sensitivity.

## Methods and Materials

### Participants

Data were collected from the initial 500 participants (323 female) from the Tulsa 1000 study (31), a naturalistic longitudinal study recruiting a community sample. Participants were between 18 and 56 years of age at the time of fMRI measurements (mean age = 35.6, standard deviation = 11.4). Participants were screened for inclusion on the basis of the Patient Health Questionnaire (PHQ-9(34)) ≥10. Exclusion criteria were positive urine drug screen; lifetime bipolar, schizophrenia spectrum, antisocial personality, or obsessive-compulsive disorders; active suicidal ideation with intent or plan; moderate to severe traumatic brain injury; severe and or unstable medical concerns; changes in psychiatric medication dose in the last 6 weeks; and fMRI contraindications. Ethical approval was obtained from Western Institutional Review Board T1000 protocol #20142082. Full exclusion criteria can be found in the parent project protocol paper (31). All participants provided written informed consent prior to participation, in accordance with the Declaration of Helsinki, and were compensated for participation. ClinicalTrials.gov identifier: #NCT02450240.

After removal of 49 participants with excessive motion in their fMRI data (i.e., participants with average Euclidian norm values across all repetition time intervals >0.3) and 235 participants who did not meet a diagnosis of either MDD or AD+MDD (see criteria below), the initial sample for the analysis included 66 participants with MDD and 150 participants with AD+MDD, defined categorically as lifetime MDD and at least one anxiety disorder according to the anxiety module of the Mini International Neuropsychiatric Interview (MINI; (35)), these include GAD, panic disorder, agoraphobia and social phobia (see **Table 1** for details). Three further participants were removed because of missing demographic data. To reduce the bias due to confounding variables, 65 participants from the MDD group were propensity matched for age, sex and education at a ratio of 2:1 with 130 participants from the MDD group using the MatchIt package in R (36). Propensity score analysis is based on the hypothesis that two patients with similar propensity scores have covariates which come from similar distributions. By selecting based on propensity scores, a new dataset is created where covariates are similar between groups (37).

**Table 1:**
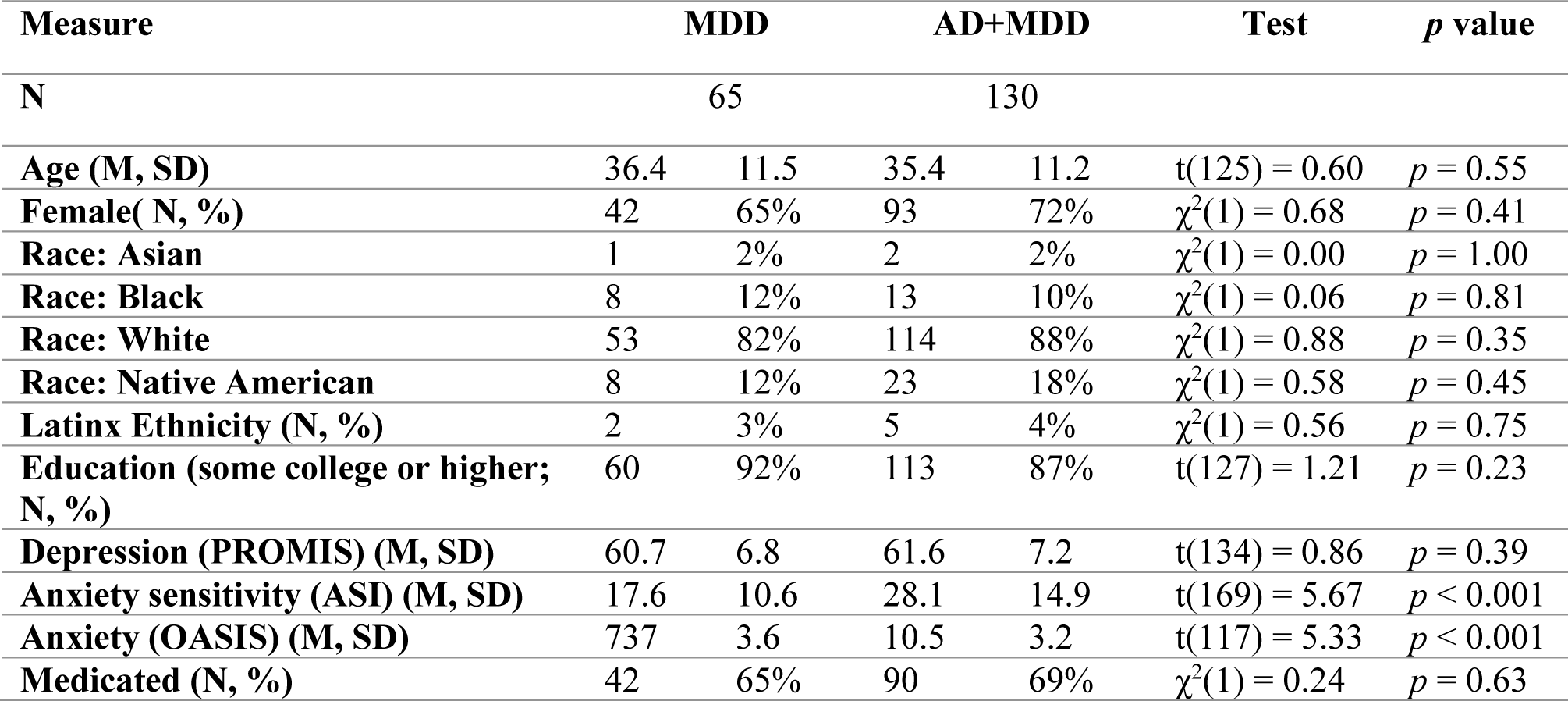
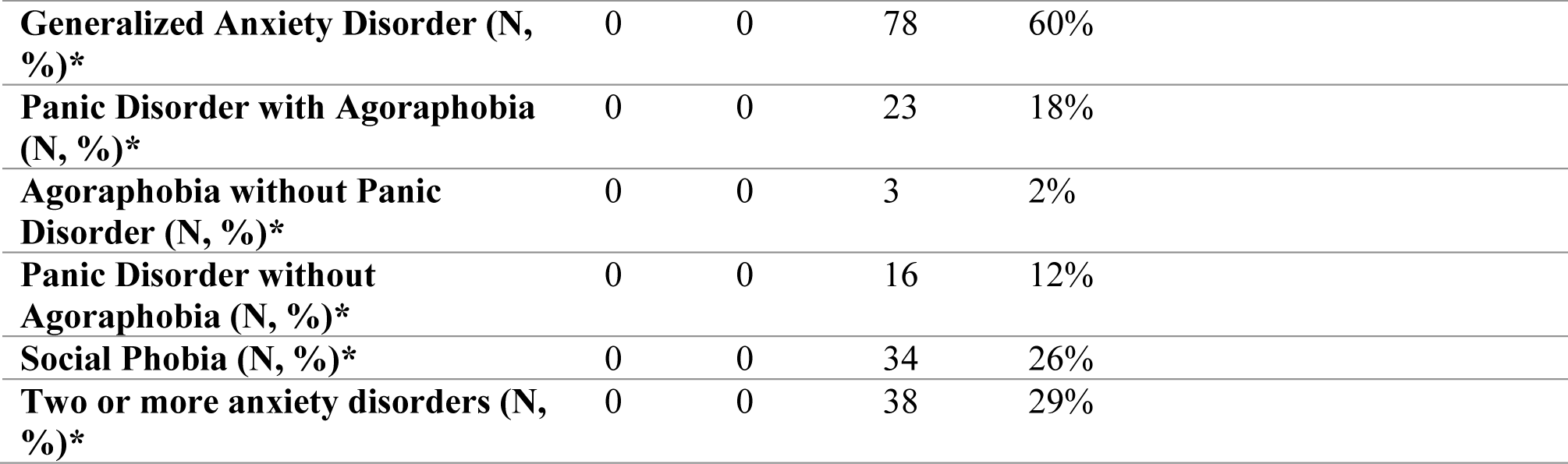
Demographic summary. Note: PROMIS: Patient-Reported Outcomes Measurement Information System, ASI: Anxiety Sensitivity Index, OASIS: Overall Anxiety and Severity and Impairment Scale.

| Measure | MDD |  | AD+MDD |  | Test | *p* value |
| --- | --- | --- | --- | --- | --- | --- |
| N | 65 |  | 130 |  |  |  |
| Age (M, SD) | 36.4 | 11.5 | 35.4 | 11.2 | $t(125) = 0.60$ | $p = 0.55$ |
| Female( N, %) | 42 | 65% | 93 | 72% | $\chi^2(1) = 0.68$ | $p = 0.41$ |
| Race: Asian | 1 | 2% | 2 | 2% | $\chi^2(1) = 0.00$ | $p = 1.00$ |
| Race: Black | 8 | 12% | 13 | 10% | $\chi^2(1) = 0.06$ | $p = 0.81$ |
| Race: White | 53 | 82% | 114 | 88% | $\chi^2(1) = 0.88$ | $p = 0.35$ |
| Race: Native American | 8 | 12% | 23 | 18% | $\chi^2(1) = 0.58$ | $p = 0.45$ |
| Latinx Ethnicity (N, %) | 2 | 3% | 5 | 4% | $\chi^2(1) = 0.56$ | $p = 0.75$ |
| Education (some college or higher; N, %) | 60 | 92% | 113 | 87% | $t(127) = 1.21$ | $p = 0.23$ |
| Depression (PROMIS) (M, SD) | 60.7 | 6.8 | 61.6 | 7.2 | $t(134) = 0.86$ | $p = 0.39$ |
| Anxiety sensitivity (ASI) (M, SD) | 17.6 | 10.6 | 28.1 | 14.9 | $t(169) = 5.67$ | $p < 0.001$ |
| Anxiety (OASIS) (M, SD) | 737 | 3.6 | 10.5 | 3.2 | $t(117) = 5.33$ | $p < 0.001$ |
| Medicated (N, %) | 42 | 65% | 90 | 69% | $\chi^2(1) = 0.24$ | $p = 0.63$ |

**Table 1: Demographic summary**
|  |  |  |  |  |
| --- | --- | --- | --- | --- |
| **Generalized Anxiety Disorder (N, %)\*** | 0 | 0 | 78 | 60% |
| **Panic Disorder with Agoraphobia (N, %)\*** | 0 | 0 | 23 | 18% |
| **Agoraphobia without Panic Disorder (N, %)\*** | 0 | 0 | 3 | 2% |
| **Panic Disorder without Agoraphobia (N, %)\*** | 0 | 0 | 16 | 12% |
| **Social Phobia (N, %)\*** | 0 | 0 | 34 | 26% |
| **Two or more anxiety disorders (N, %)\*** | 0 | 0 | 38 | 29% |

### Procedure

General procedures included a clinical interview session, a neuroimaging session, and a behavioral session, completed within two weeks on average. Although the parent project (i.e., T1000) consisted of a broader range of protocols, only details relevant to the current study are presented here. See protocol paper (31) for full details. Study staff administered the MINI clinical interview and collected self-reported information on demographics.

### The fear learning task

The fear conditioning/extinction (FC/FE) paradigm (**Fig. 1**) was previously described in (38). It was based on Pavlovian conditioning and the task previously used in neuroimaging studies of individual differences in fear learning (39–41). The stimuli consisted of two neutral, non-social, abstract images as conditioned stimuli (CS), presented for 1.5 s at a time. The images designated as CS+ (paired with the unconditioned stimulus (US) during conditioning) and CS-(never paired with the US) were counter-balanced across participants. The US consisted of a loud scream beginning 500 ms after CS+ onset, lasting approximately 1 s, and presented at 108-120DBs with participants wearing silicone ear plugs providing 22DBs of attenuation.

**Figure 1:**
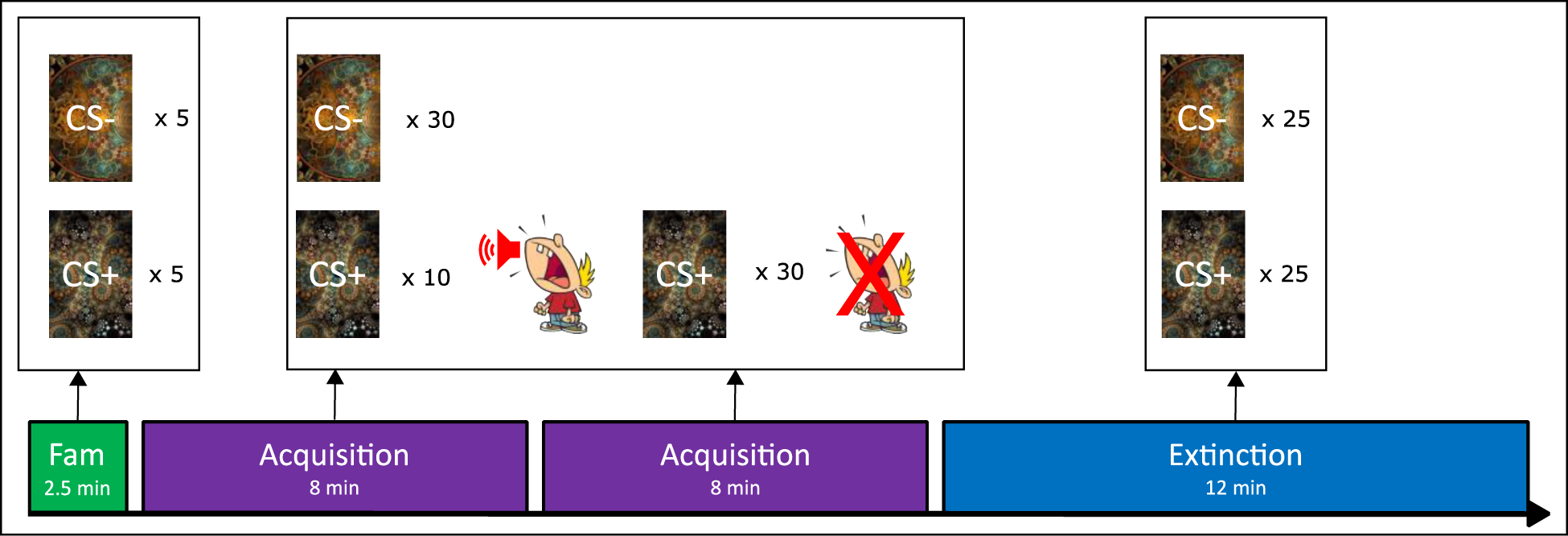
Pavlovian Fear Conditioning Paradigm. During the task, participants viewed two neutral, non-social, abstract images, one of which was paired with a loud scream. The task consisted of three consecutive phases: familiarization (Fam), acquisition, and extinction. Following each run, participants rated their levels of anxiety, arousal, and image valence. Images were counterbalanced across participants. Trials were presented in a fixed, pseudo-randomized order. CS+ = conditioned stimulus that was paired with the scream on some trials during acquisition; CS-= conditioned stimulus that not paired with the scream.

**Figure 2.**
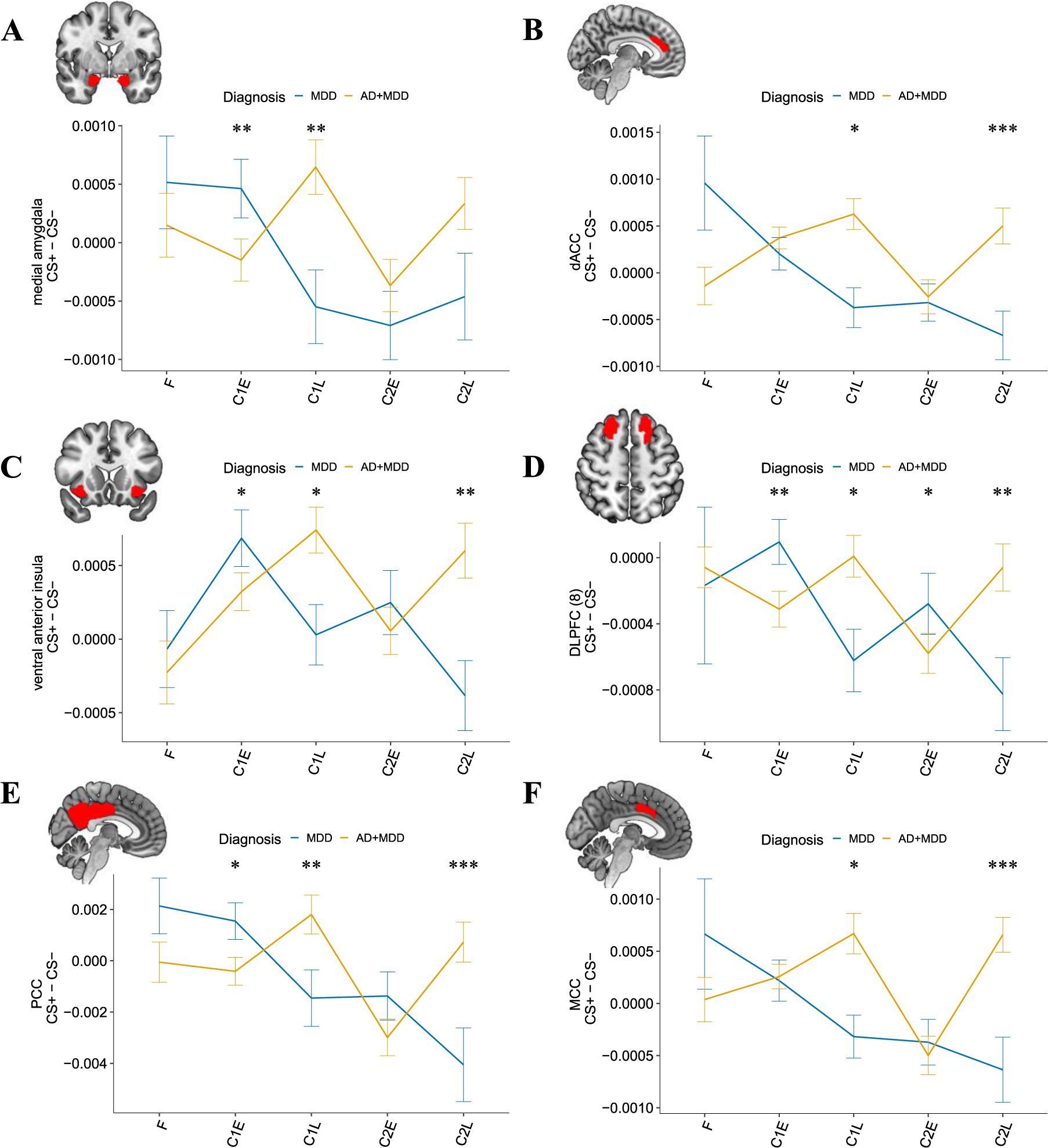
Group differences in neural fear conditioning responses. Group differences in percent signal change comparing fear acquisition to familiarization in the CS+ vs. CS-contrast in A) medial amygdala; B) dorsal Anterior Cingulate Cortex (dACC); C) ventral anterior insula; D) dorsolateral prefrontal cortex (DLPFC); E) Posterior Cingulate Cortex (PCC); F) Midcingulate Cortex (MCC). Acquisition runs are split into early and late.

The task consisted of three phases: familiarization, fear conditioning, and fear extinction. The familiarization phase (lasting 2.5 min) involved five presentations of each CS with no instances of the US. Next, the conditioning phase involved 30 presentations of the CS- and 40 presentations of the CS+ (10 with (CS+paired) and 30 without (CS+unpaired) the US), across two functional runs of eight minutes each.

### fMRI Data Processing and Analysis

Imaging data were acquired at the Laureate Institute for Brain Research using two identical GE MR750 3T scanners and an 8-channel phased-array coil. The following parameters were used for all EPI data: TR/TE=2000/27ms, FOV/slice = 240/2.9mm, 128×128 matrix, 39 axial slices, and varied numbers of TRs depending on the run (familiarization=79, conditioning 1/2=260 each, and extinction=368). For normalization to standard space, a high-resolution Magnetization-Prepared Rapid Acquisition with Gradient Echo sequence was also acquired with the following parameters: TR/TE=5/2.012 ms, FOV/slice = 240×192/0.9mm, and 186 axial slices.

The AFNI (http://afni.nimh.nih.gov) software package was used for all first-level neuroimaging data analyses (42). Processing steps included: removal of the first three volumes, despiking, slice timing correction, co-registration with anatomical volumes, motion correction, 4mm full width at half maximum Gaussian smoothing, scaling to percent signal change, normalization to MNI space using an affine transformation and resampling to 2mm isometric voxels. Censoring was applied at the regression step using a Euclidean norm motion threshold of 0.3.

We focused our analysis on the acquisition/conditioning phases compared to familiarization. Previous research has implicated several regions in conditioning (25, 27, 43–46). Driven by these past findings, we explored task effects from both hemispheres of the following probabilistic cytoarchitectonic segmentations defined by the Brainnetome atlas (BA) (47): amygdala (lateral and medial), anterior mid-cingulate cortex (aMCC; BA caudodorsal 24), dorsal anterior cingulate cortex (ACC; BA pregenual 32), rostral hippocampus (rostral and caudal), anterior insula (ventral and dorsal), ventromedial prefrontal cortex (vmPFC), posterior cingulate cortex (PCC; composed of BA ROIs left: 153, 175, 181, 185; right: 154, 176, 182, 186), and two regions of DLPFC (area 6 and area 8). For the main analysis, the average percent signal change (PSC) for the contrast of CS+ vs. CS-were extracted from these ROIs for five different time periods: familiarization (F), Conditioning Run 1, Early (C1E) and Late (C1L), Conditioning Run 2 Early (C2E) and Late (C2L) and subjected to lmer (model: beta ∼ group * time + sex + age + medication + motion (1|id)), as well as follow-up analyses using the lmertest package in R, with type III analysis of variance significance tests of interactions. Follow up analyses additionally examined CS+ and CS-separately and on a trial-by-trial basis. A Bonferroni correction was applied for number of ROIs (12), corrected *p* values are reported for the omnibus tests. Whole brain analyses and extinction analyses are reported in the Supplemental Information.

## Results

### Demographic and clinical data

The groups did not differ on demographics such as age, sex, and education, and had similar levels of medication (all *p* < 0.23, see **Table 1**). Groups also did not differ on severity of depression (Patient reported outcomes measurement information system; PROMIS(48); *p* = 0.39) but, crucially, had significantly different levels of trait self-report anxiety sensitivity (Anxiety sensitivity Index; ASI (49)) and state anxiety severity and impairment (Overall anxiety severity impairment scale; OASIS (50)) (both *p* < 0.001).

### Neuroimaging results

For CS+ vs. CS-there were significant main effects of *Time* (conditioning phases vs. F) in the PCC, rostral hippocampus and vmPFC, (all *p_corr_* < 0.009) indicating that the task elicited changes in neural activation overall.

For CS+ vs. CS-there were significant *Group* x *Time* interactions in the medial amygdala, dorsal ACC, ventral anterior insula, DLPFC (area 8), MCC and PCC (all *p_corr_* < 0.04). Overall, there is a pattern for increased initial activation in MDD in four of these regions which decreased over the course of the conditioning runs, whereas the AD+MDD group show increased sustained neural reactivity in all these regions over time within each conditioning run.

An examination of the contrasts (*Group:* AD+MDD vs. MDD; *Time:* 4 conditioning phases vs. F) showed that for the medial amygdala, ventral anterior insula, DLPFC (area 8), and the PCC, the MDD group had greater activation in C1E (vs. F) compared to the AD+MDD (all *p* < 0.023) group whereas the AD+MDD group showed greater activation in all these regions as well as the dorsal ACC and MCC during C1L (vs. F) (all *p* < 0.034) compared to the MDD group. Follow-up analysis examining stimulus type separately showed these regional differences to be driven by higher responses to the CS+ in AD+MDD vs. MDD during C1L (vs. F) (all *p* < 0.007). Additionally, lower responses to CS-in AD+MDD vs. MDD were found in the dorsal ACC and MCC during C1L (vs. F) (all *p* < 0.03).

For the DLPFC (area 8), an examination of contrasts revealed greater activation in the MDD group (compared to AD+MDD) during C2E (vs. F) (*p* = 0.034) whereas greater activation in the AD+MDD group was seen in C2L (vs. F) in the DLPFC (area 8) as well as dorsal ACC, ventral anterior insula, MCC, and PCC (all *p* < 0.005). Follow-up analysis examining stimulus type showed this to be driven by higher responses to CS+ in AD+MDD vs. MDD during C2L (vs. F) in the dorsal ACC, ventral anterior insula, MCC, and PCC (all *p* < 0.02) and lower responses to CS-in AD+MDD vs. MDD in the same regions except for the ventral anterior insula (all *p* < 0.008). For summary of all *Group* x *Time* statistics, see Table 2.

**Table 2:**
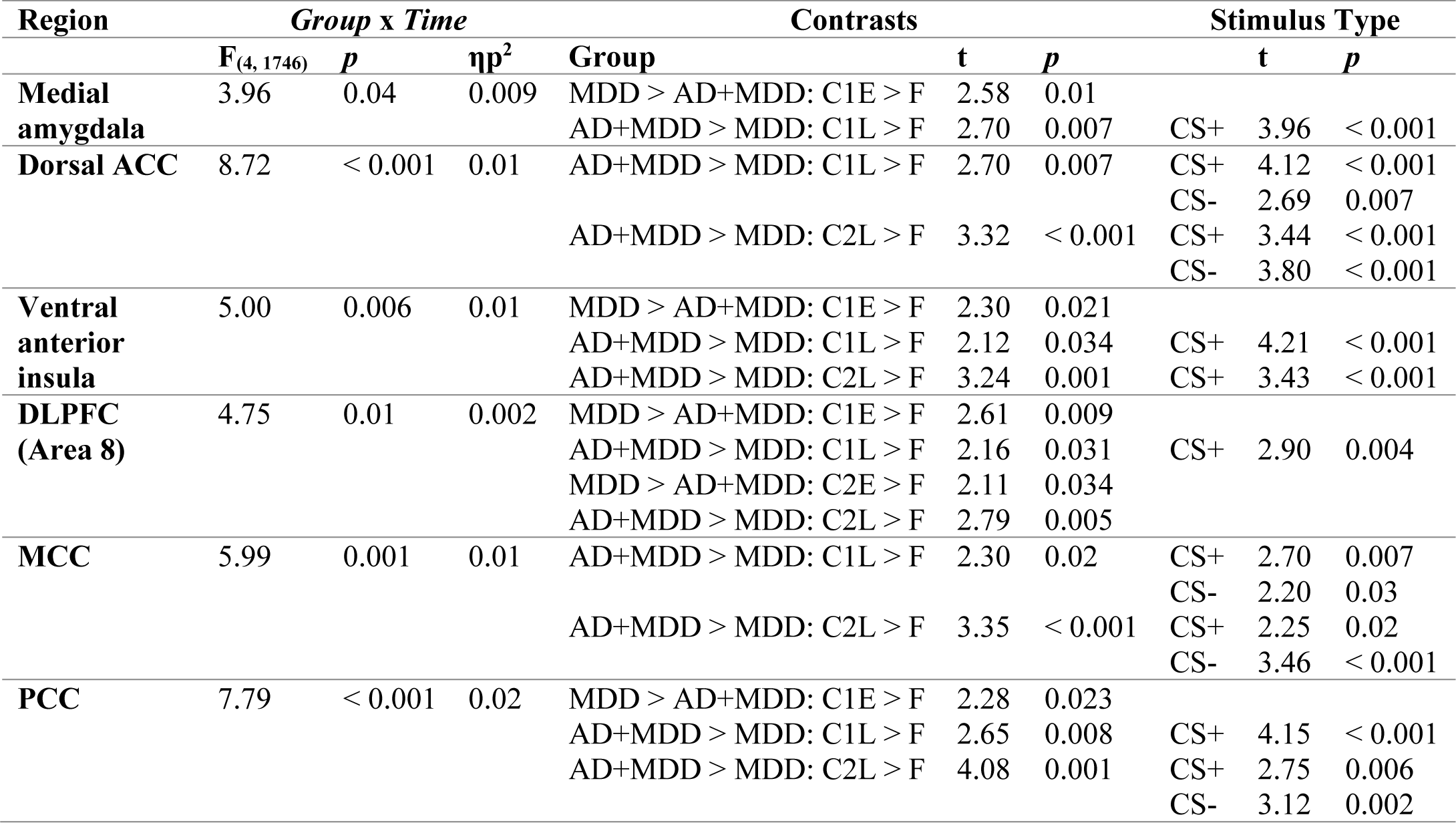
Results summary. Note: Dorsal ACC: dorsal anterior cingulate cortex, DLPFC: dorsolateral prefrontal cortex, PCC: posterior cingulate cortex, MCC: midcingulate cortex, F: familiarization, C1E: conditioning run 1 early, C1L: conditioning run 1 late, C2E: conditioning run 2 early, conditioning run 2 late.

No main effects of *Group*, or *Time* or any *Group* x *Time* interactions survived correction for the lateral amygdala, caudal hippocampus, dorsal anterior insula, or DLPFC (area 6) (all *p_corr_* > 0.06). No significant main effects of *Group, Sex, Age, or Laterality* were seen for any ROI. Whole brain analyses are reported in the supplement.

## Discussion

The primary objective of this study was to investigate whether individuals with comorbid AD and MDD exhibit a distinct profile of exaggerated defense-related processes, specifically in threat sensitivity, compared to those with MDD alone. This was assessed examining activation patterns during fear conditioning in key regions in the threat circuitry. Our results revealed significant *Group* x *Time* interactions in several regions, including medial amygdala, dACC, ventral anterior insula, MCC and PCC. Notably, the MDD group showed increased initial activation in four of these regions that decreased over the course of the conditioning runs. In contrast, the AD+MDD group exhibited sustained neural reactivity in all these regions throughout each conditioning run. Follow-up analyses further substantiated these differences, as effects were consistently driven by the CS+ condition, and can thus be interpreted as fear signaling, rather than safety signaling. Based on these findings, we can preliminarily conclude that AD+MDD present a unique neural profile of exaggerated threat sensitivity compared to MDD alone. This is evidenced by sustained activation in key regions of the threat circuitry during fear conditioning, contrasting with the more transient and later decreased activation observed in the MDD group. Therefore, we underscore the necessity for developing targeted pharmacological and psychological interventions that directly modulate the sustained neural reactivity observed in the AD+MDD population.

The AD+MDD group showed increased DLPFC activation, which was also driven by the CS+ condition, perhaps reflecting greater demand from hyperactive limbic structures for regulation during fear acquisition. As with the fear circuitry, after initial activation at the beginning of conditioning, the MDD group also showed overall blunted regulatory responses to conditioning. This also suggests that fear conditioning processes are different in those with AD+MDD compared to MDD and may represent a useful treatment target.

Prior studies comparing MDD and AD+MDD are rare, particularly in fear conditioning and neuroimaging. Thus, to our knowledge this is the first study that examines differential neural correlates of fear conditioning in these two groups. A recent meta-analysis of fear conditioning in AD including 900 AD patients and 1200 healthy comparisons (51) suggested overgeneralization of fear responses to safety cues but mixed findings otherwise, citing heterogeneity of the included studies. Fear conditioning in MDD is less well studied than AD, with a limited number of studies reporting enhanced (52), impaired (53) and similar (54) levels of fear acquisition (measured with skin conductance and eyeblink startle) to healthy comparisons. We speculate that effects of comorbid AD+MDD may be one source of heterogeneity that could be masking the potentially opposing effects of MDD and AD in these studies.

We previously showed that AD+MDD was associated with greater threat sensitivity measured with emotionally modulated eyeblink startle response (32) and interoceptive/nociceptive fearfulness/reactivity (33) compared to those with MDD alone. These findings start to elucidate the neural mechanisms that may underlie threat sensitivity in AD+MDD. The interoceptive and exteroceptive features of threat sensitivity have a different putative neural basis (55, 56). Eyeblink startle response has been directly linked to amygdala activation (57) and dACC and insula are involved in autonomic-interoceptive integration (58, 59). These regions all showed group differences in the present study. Increased response in AD+MDD compared to a more blunted response in MDD alone at these multiple levels of measurement provides evidence that these two subtypes have different process dysfunction and potentially should not be merged as a single group for experimental studies or clinical trials. This also offers a potential explanation for prior mixed findings, with group averages being conflated by opposing hyperarousal and blunting.

Like the different characteristics of “fear” and “distress” disorders within the wide-ranging category of ADs, threat sensitivity is a broad term that necessitates a more precise conceptualization. There is some debate about how the constituent parts of threat processing should be defined but one popular approach (adopted by the NIMH research domain criteria) focuses on the imminence of the threat. Under this, acute threat is near (representing “fear”) and potential threat is more distant (representing “anxiety”). Pavlovian fear conditioning does not separate out these phenomena neatly, as there is some aspect of acute and potential threat during conditioning. This dual-model theory of fear conditioning in humans (60) describes two complementary defensive systems: a reflexive lower-order system independent of conscious awareness and a higher-order cognitive system associated with conscious awareness of danger and expectation. These processes map onto “fear” and “distress” disorders respectively, making studies of fear conditioning relevant across all the ADs. However, future studies should employ paradigms that seek to better characterize threat sensitivity in terms of acute and potential threat (16, 61) so that more precise mechanisms may be illuminated.

If AD+MDD and MDD do have the proposed process function differences, there are possible implications in terms of treatment for both MDD and AD+MDD. For example, the understanding of potential target processes (e.g. fear) and associated circuits (e.g. dACC, amygdala) could inform neuromodulatory interventions for AD+MDD. A recent translational study was able to enhance extinction learning with paired transcranial magnetic stimulation (TMS) to frontal regions functionally connected with vmPFC (62). Neurofeedback paradigms could also be informed by validation of these target processes and circuits in AD+MDD, such as a recently developed amygdala neurofeedback intervention for stress resilience (63). In terms of pharmacological treatments, novel agents (e.g. neuroactive steroids such as Brexanolone (64)) that target neurotransmitters (e.g. GABA) typically modulated by more traditional anxiolytic agents (e.g. lorazepam) could potentially have better efficacy in AD+MDD. Finally, for individuals with MDD alone, cognitive therapy focused on engaging with threat or any type of salience in their environment might be beneficial (e.g., behavioral activation) whereas for AD+MDD, efforts may be more usefully focused on acceptance of threat (e.g., acceptance-based strategies during exposure exercises).

The observed differences in neural reactivity between the AD+MDD and MDD groups during fear conditioning raise several intriguing questions about the underlying neurobiological mechanisms responsible for these distinct profiles. First, future studies should aim to explore a range of potential factors, including using magnetic resonance spectroscopy (MRS) to examine neurotransmitter dysregulation that are critical for neural plasticity, such as imbalances in excitatory (glutamatergic) and inhibitory (GABAergic) neurotransmission. Second, these findings could be the result of dysregulation of the hypothalamic-pituitary-adrenal axis and cortisol levels. Third, on a behavioral level cognitive bias, such as attentional and interpretational biases, and their neural underpinnings are another promising area for exploration. On a neural level, impaired top-down regulation from cognitive control centers like the DLPFC could be a critical factor and altered functional connectivity between limbic and prefrontal regions may contribute to the observed patterns and should be examined using advanced neuroimaging techniques. Understanding these specific neurobiological processes will not only deepen our understanding of the complex interplay between anxiety and depression but also inform the development of targeted pharmacological and cognitive-behavioral interventions for this particularly challenging clinical population.

Limitations of the study include the cross-sectional design, the use of a scream US rather than shock (65) and half the number of MDD as AD+MDD participants. Nevertheless, to our knowledge this study reports the largest sample used to compare AD+MDD and MDD individuals on an fMRI fear conditioning task.

In conclusion, this study provides compelling evidence that individuals with comorbid AD+MDD show a distinct neurobiological profile of exaggerated threat sensitivity compared to those with MDD alone, underscoring the need for targeted interventions. These findings align with and extend previous work, shedding light on the complex interplay between anxiety and depression at the neural level. They also highlight the importance of not merging AD+MDD and MDD into a single group for experimental studies or clinical trials, given their distinct neural and behavioral profiles. Future research should focus on a multi-faceted exploration of underlying neurobiological mechanisms, ranging from neurotransmitter dysregulation to cognitive biases and their neural substrates. Such comprehensive investigations will not only refine our understanding of these disorders but also pave the way for the development of targeted pharmacological and psychological interventions, potentially including neuromodulatory techniques and neurofeedback paradigms, that are specifically tailored to address the sustained neural reactivity observed in the AD+MDD population.

## Acknowledgements

This work has been supported in part by The William K. Warren Foundation and the National Institute of General Medical Sciences Center Grant Award Number 1P20GM121312. The content is solely the responsibility of the authors and does not necessarily represent the official views of the National Institutes of Health.

The ClinicalTrials.gov identifier for the clinical protocol associated with data published in the current paper is NCT02450240, “Latent Structure of Multi-level Assessments and Predictors of Outcomes in Psychiatric Disorders”.

The authors thank all the research participants and wish to acknowledge the contributions of Tulsa 1000 Investigators towards the collecting and organizing of data.

Maria Ironside, Rayus Kuplicki, Jennifer Stewart, Robin Aupperle, and Martin Paulus receive funding from the National Institute of General Medical Sciences (NIGMS) center grant P20GM121312; Maria Ironside has additional funding from National Institute of Mental Health (NIMH; R01MH132565). Sahib Khalsa has grant funding from the NIMH (K23MH112949, R01MH127225); Robin Aupperle has additional grant funding from NIMH (K23MH108707; R01MH123691); Jennifer Stewart has additional grant funding from the National Institute of Drug Abuse (NIDA) (R01DA050677), Rayus Kuplicki has additional funding from NIDA (R01DA050677); and Martin Paulus has additional grant funding from the National Institute of Drug Abuse (U01DA041089, R01DA050677).

