## Supplemental Information for "The unique face of anxious depression: Increased sustained threat circuitry response during fear acquisition"

### Supplemental Methods

***Fear conditioning task:*** The 25% reinforcement rate is consistent with other fMRI studies of individual differences in fear conditioning (1, 2). CS+paired trials were modeled in the individual-level deconvolution analysis but omitted from the group-level CS+ analysis to not confound processing of the CS+ with the reactivity to the US. This also allowed for an equal number of trials in the analysis. Finally, the extinction phase (lasting 12 min) followed immediately after conditioning and involved 25 presentations of each CS with no US reinforcement. Participants rated their anxiety level using a Likert scale ("On a scale from 0 = minimum anxiety to 100 = maximum anxiety, how anxious do you feel when you see this image?"), as well as valence ("How happy or unhappy does this image make you feel?") and arousal ("How calm or excited does this image make you feel?") levels using the Self-Assessment Manikin (3) to each CS after each functional run. Trials were presented in a fixed, pseudo-randomized order, constrained so that no more than two identical trials occurred in a row.

***Whole brain fMRI analyses:*** AFNI's 3dMVM was used to fit voxel-wise multivariate models of the CS+ vs. CS- contrast as the dependent variable, including fixed effects for *group* (MDD, AD+MDD), *timepoint* (F, C1E, C1L, C2E, C2L, EE, EL), and the *group* x *time* interaction. Each conditioning run (C1E, C1L, C2E, C2L) was contrasted with F and the two halves of the extinction run (EE, EL) were contrasted with C2L. The smoothness of the group level error terms was estimated with AFNI's 3dFWHMx using the spatial autocorrelation function (acf) (4) and used with 3dClustSim to produce cluster size thresholds controlling the family-wise error rate (-acf a, b, c parameters: 0.56, 3.22, 9.14). Significance criterion for the whole-brain analysis was set at a corrected rate of $p < 0.05$ (cluster size ≥83 voxels) and thresholded per-voxel at $p < 0.001$.

**Supplemental Results:**

***Whole brain fMRI results:*** For the contrast of C2E vs F there were 10 negative signed clusters that survived correction (see table S1) and for the contrast of EL vs C1L there was increased activation in the dorsolateral prefrontal cortex (DLPFC) (XYZ: +39, -21, +45, K=102) and the right angular gyrus (XYZ: -49, +63, +41, K=128). There were no significant *group* x *time* interactions for any of the conditioning runs compared to F. In C1L the AD+MDD group showed higher activation in the bilateral superior medial gyrus (XYZ: -9, -59, +35, K=219, see Fig. S1A) and in an area overlapping the right DLPFC and frontal operculum (XYZ: -49, -15, +29, K=99, see Fig. S1B) compared to the MDD group. For the contrast of EL vs C2L, MDD group showed increased activation compared to the AD+MDD group in the left inferior frontal gyrus (XYZ: -49, -23, -23, K=83, see Fig S1C). However, the same session extinction protocol used in the task and resultant baseline differences (from using C2L as baseline for extinction) preclude interpretation of this group effect.

| **Significant clusters for C2E vs F**<br>**Region** | **Peak co-ordinates in MNI space** | | | **Voxels** |
|---|---|---|---|---|
| Left cuneus/ superior occipital cortex | +19 | +83 | +43 | 796 |
| Right superior temporal gyrus | -51 | +29 | 13 | 593 |
| Right precentral gyrus | -35 | +27 | +65 | 460 |
| Left Superior temporal gyrus | +59 | +3 | -3 | 322 |
| Mid-cingulate cortex | -1 | +25 | +57 | 266 |
| Left precentral gyrus | +25 | +27 | +71 | 260 |
| Left lingual gyrus | +9 | +77 | -7 | 147 |
| Right lingual gyrus | -7 | +51 | +3 | 129 |
| Left lingual gyrus | +11 | +59 | -5 | 112 |
| Right postcentral gyrus | -61 | +17 | +43 | 84 |

**Table S1: Significant negative clusters for the contrast C2E vs F**

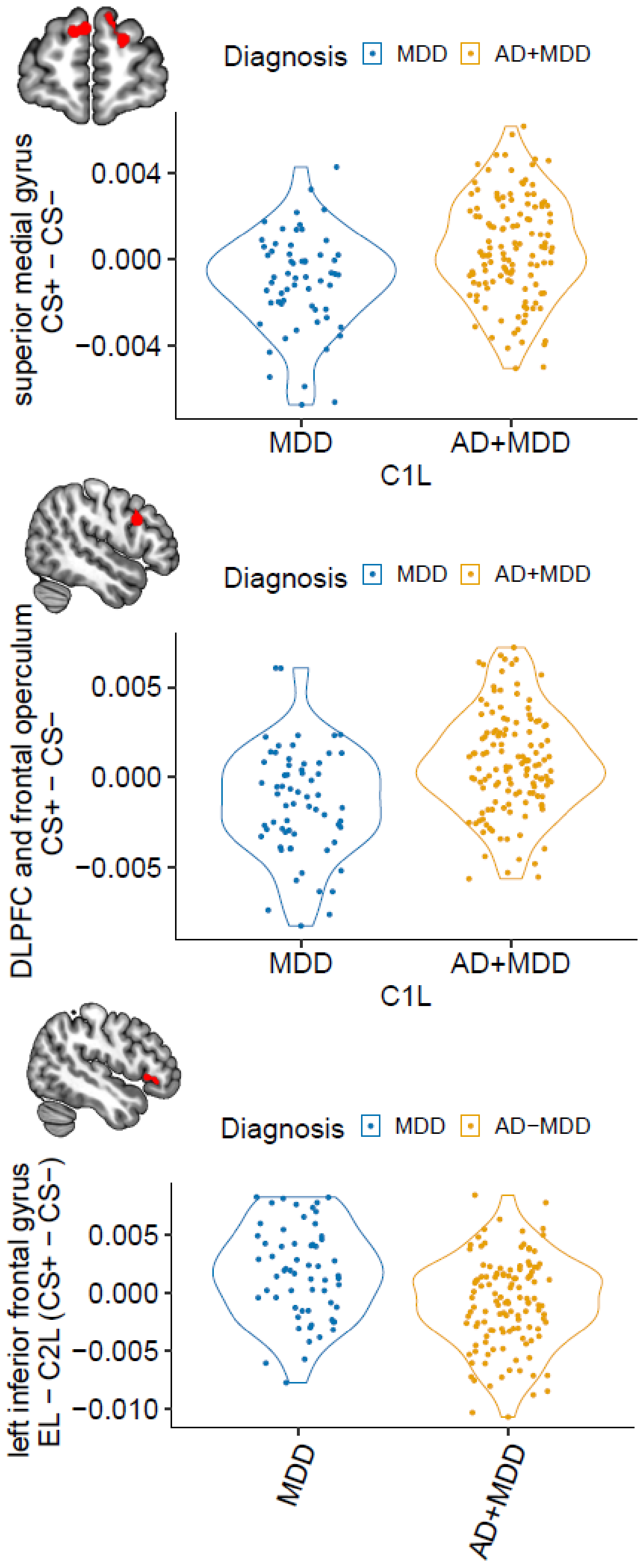


**Suplemental Figure 1: Whole brain findings**
